# Population-level age effects on the white matter structure subserving cognitive flexibility in the human brain

**DOI:** 10.1101/2024.08.30.610526

**Authors:** Tatiana Wolfe, Alexandra Gassel, Maegan L. Calvert, Lee Isaac, G. Andrew James, Timothy R. Kosick, Clint D. Kilts

## Abstract

Cognitive flexibility, a mental process crucial for adaptive behavior, involves multi-scale functioning across several neuronal organization levels. While the neural underpinnings of flexibility have been studied for decades, limited knowledge exists about the structure and age-related differentiation of the white matter subserving brain regions implicated in cognitive flexibility. This study investigated the population-level relationship between cognitive flexibility and properties of white matter across two periods of adulthood, aiming to discern how these associations vary over different life stages and brain tracts. We propose a novel framework to study age effects in brain structure-function associations. First, a meta-analysis was conducted to identify neural regions associated with cognitive flexibility. Next, the white matter projections of these neural regions were traced through the Human Connectome Project tractography template to identify the white matter structure associated with cognitive flexibility. Then, a cohort analysis was performed to characterize myelin-related macromolecular features using a subset of the UK Biobank magnetic resonance imaging (MRI) data, which has a companion functional/behavioral dataset. We found that (1) the wiring of cognitive flexibility is defined by a subset of brain tracts, which present undifferentiated features early in adulthood and significantly differentiated types in later life. (2) These MRI-derived properties are correlated with individual subprocesses of cognition, which are closely related to cognitive flexibility function. (3) In late life, myelin-related homogeneity of specific white matter tracts implicated in cognitive flexibility declines with age, a phenomenon not observed in early life. Our findings support the age-related differentiation of white matter tracts implicated in cognitive flexibility as a natural substrate of adaptive cognitive function.

**Significance Statement:** Cognitive flexibility function facilitates adaptation to environmental demands. Brain changes affecting structural organization during the lifespan are theorized to impact cognitive flexibility. This study characterizes how the brain’s connectivity is correlated with cognitive flexibility function throughout adulthood. By analyzing myelin-related properties of white matter, this study found that certain parts of the brain’s wiring related to cognitive flexibility become more differentiated with advanced age. These age-related features appear as a natural characteristic of the human brain that may impact specific aspects of adaptive thinking, like shifting between tasks or updating information.

## Introduction

Cognitive flexibility is an inherent ability that is important for human adaptive functioning and behavior.^1^ This mental process is acquired during development and continues to evolve over the adult lifespan.^2,3^ It plays a crucial role in various aspects of life, including coordination with other executive functions and behaviors.^4^ Impairments in cognitive flexibility are common among several neurological and neuropsychiatric disorders, including anxiety and dementia;^6,7^ and maintaining cognitive flexibility in later life may help mitigate cognitive decline associated with late-life aging. Thus, to understand the intrinsic properties of the neural mechanisms of human adaptive thinking and behavior, it is important to elucidate the effectors relevant to cognitive flexibility function during adulthood.

Cognitive flexibility involves brain-wide functioning at several neural organization levels, including degrees of regional neural liberality (e.g., the ability to switch processing assignment in synchronized pace with the functional network)^7,8^ and shifts in network-wide properties of brain function (e.g., connectivity or eccentricity).^4,9,10^ Over the course of natural aging, neural organization changes regionally through synaptic pruning,^11^ cortical myelination,^12^ and related activity-dependent tailoring mechanisms,^13^ as well as at brain-wide network levels through frequency-dependent integration (e.g., between-network connectivity).^14^ Visible changes in the interconnecting white matter accompany these functional transitions.^15^ The trajectory of white matter maturation over the lifespan varies among different brain tracts and neuropils,^16^ which modifies how brain regions decentralize functional processing while increasing regional specialization.^13^ This process has direct implications for the topography of functional networks and, consequently, cognitive functioning.

Imaging studies that measure the number of white matter fiber tracts connecting brain regions have shown that advanced age affects white matter tracts differently, with prefrontal tracts showing greatest age-related thinning in humans.^17^ These studies frequently report age-related changes in white matter as detrimental alterations to tissue structure. On the other hand, histopathological studies that reveal macromolecular lipid density through staining—a quantity related to tract myelination—have further noted that deep white matter and its projections into the cerebral cortex show highly differentiated arrangements with age that are not necessarily reflective of pathology.^18^ Sherin and Bartzokis have summarized that, in older adults, white matter fibers within bundles appear less dense and show robust selective myelination along their projections and greater numbers of myelinated connections within the gyri. In younger adults, fiber myelination is rather abundant and largely undifferentiated within the bundles. These cellular structural characteristics of brain white matter demonstrate a level of age-related adaptation in brain tissues.

Models of aging-related differentiation of white matter have been proposed.^13^ In an influential review, Bonetto et al. outlined the processes of activity-dependent myelination as a natural adaptive brain process that modifies the white matter in response to function—rather than functional changes being a consequence of (de)myelination.

In this work, we computed features of white matter derived from magnetic resonance imaging (MRI) measures of macromolecular tissue properties (rather than diffusion), and investigated their association with cognitive flexibility and age. We propose a novel method to derive the population-level structure of cognitive flexibility and investigate age effects (**Fig. 1**). This framework aims to investigate aspects of white matter differentiation by comparing MRI features derived from T1-weighted (Tw) and T2-fluid-attenuated inversion recovery (T2-FLAIR) data to functional scores dimensionally. We focus on a brain representation of cognitive flexibility that is not explicitly associated with any pathology; thus, the structural and functional variability encompassed in the analyses is reflective of natural aging processes. Specifically, we define the white matter tracts that are particularly implicated in cognitive flexibility function and test three related hypotheses: (1) that an intrinsic association exists between cognitive flexibility capacity and the homogeneity of the white matter implicated in cognitive flexibility function; (2) that this relationship differs between early and later adulthood; and (3) that these associations exist in either life period in a tract-selective manner.

**Figure 1.**
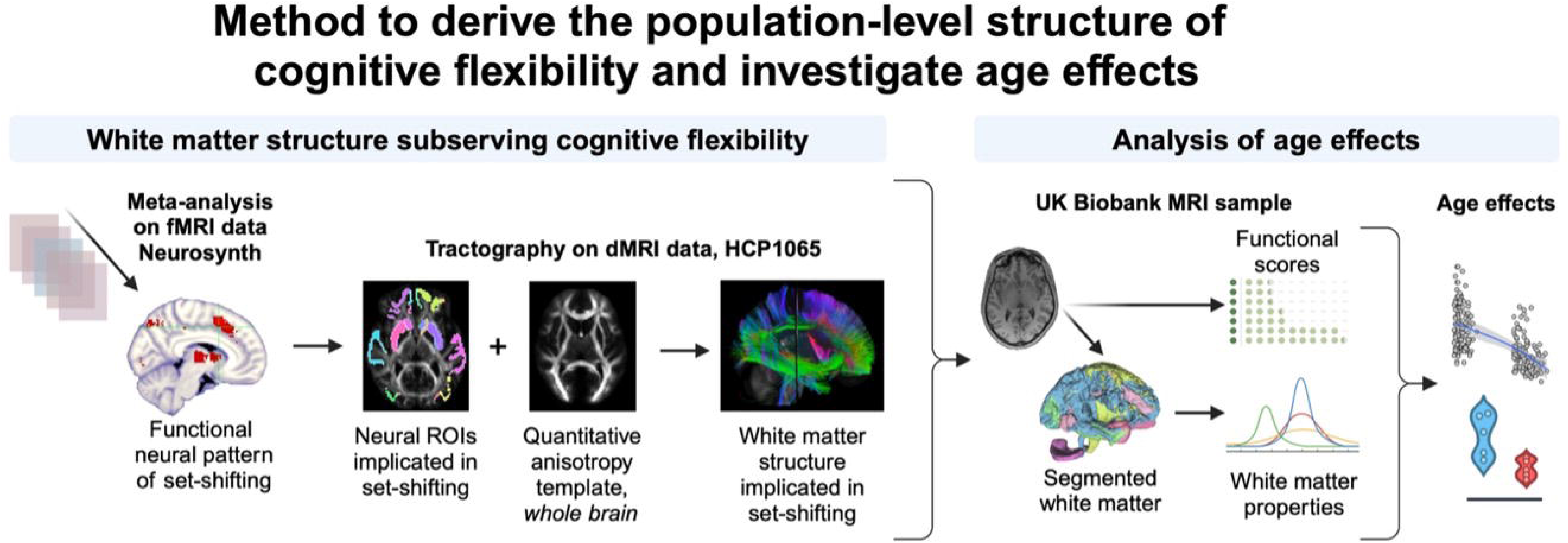
Methodological approach to investigate age effects in brain white matter structures implicated in cognitive flexibility function. First, the functional neural pattern was identified by meta-analysis using Neurosynth (left) as a collection of imaging voxels reportedly implicated in set-shifting function in human experiments. These voxels derived from functional magnetic resonance imaging (fMRI) data were then considered neural regions of interest (ROIs), which served as input for diffusion-based tractography (center). The Human Connectome Project (HCP) 1065 template of diffusion MRI (dMRI) quantitative anisotropy was used to trace the white matter structure interconnecting set-shifting ROIs. Tracts were anatomically identified using cluster recognition in DSI-studio. The collection of tracts identified as implicated in the set-shifting function composed the structure of cognitive flexibility referred to in this work. Next, a comparative cohort analysis was conducted using a subset of the UK Biobank imaging data (n=301), which has a companion table of cognitive and behavioral scores for set-shifting and related functions. Following the segmentation of white matter bundles at the individual level among the sample, two macrostructural properties of the white matter were computed (right): the mean intensity in standardized T1w MRI (*m*) and the kurtosis of the distribution of the T1w/T2-FLAIR ratio (*k*). These properties were computed per subject, for each white matter bundle individually and for the whole aggregate. Lastly, the effects of age were quantified by modeling the relationship between the image-derived macrostructural properties of white matter, functional scores, and age or age group.

## Materials and Methods

### Brain Structure Implicated in Cognitive Flexibility

#### Meta-analysis

Neurosynth (Neurosynth.org) is a platform designed for automated, large-scale meta-analysis, which synthesizes brain map data from thousands of studies banked in the Neurosynth database. Our results reflect a limited search conducted early in 2024 when the database was reported to encompass data from 14,371 studies and 1,334 searchable terms.^19^ The database is naturally limited by the existing literature archive. To identify neuroimaging results from brain regions that are reported to be active during dimensional set-shifting tasks, we used the term ‘switching,’ which was consistently used to describe dimensional set-shifting neuroimaging studies that we aimed to include in our analysis. Switching—or set-shifting—is a component of the broader functional class of executive functioning, which also involves inhibitory control and information-updating abilities.^1^ Here, the single term ‘switching’ was used aiming to evoke studies reporting brain function associated primarily with set-shifting.

Two brain maps and the under-layered anatomical volume (i.e., Montreal Neurological Institute template) were obtained from the search in Neurosynth, including the uniformity test map and the association test map. Both maps are corrected for false discovery rate, with an expected rate of 0.01 or less. The 1 mm^3^ isotropic-voxel uniformity map consists of all brain voxels reported in neuroimaging studies to be correlated with dimensional switching abilities. The association test map consists of voxels reported as also being correlated with other abilities, for example, the term ‘language’ in a language-switching task. The resulting specificity (S) of the term search where the ratio between the total number of voxels present in the association map (N_a_) and the total number of voxels present in the uniformity test map (N_u_) was computed as:

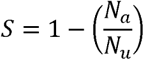

An arbitrary threshold of effect sizes >2 was applied to the association test map to remove spurious associations while estimating the sensitivity of the term search. This threshold is set in Neurosynth to modify the association test map only (not the neural activations from the meta-analysis). Voxels that are implicated in the term search may also be implicated in other searchable terms (e.g., learning). The effect size threshold controls the relative magnitude of association between a voxel and the term ‘switching,’ specifically, in comparison with the magnitude of association between the same voxel and other search terms in the database. A threshold of effect sizes >2 was chosen to estimate the specificity of the search in terms of voxels that are at least two times more likely to associate with the search term.

A list of Montreal Neurological Institute coordinates was extracted for all voxels in the meta-analysis uniformity map using MRIcron (www.nitrc.org/projects/mricron). A systematic search, including four distinct atlases of the human brain, was conducted to identify cortical and subcortical regions corresponding to the extracted coordinates. Brainnectome ((atlas.brainnetome.org), Atlasing of the Basal Ganglia ‘ATAG_basal_ganglia’ (www.nitrc.org/projects/atag), FreeSurfer Desikan-Killiani cortical parcellation ‘FreeSurfer_DKT_cortical’ (surfer.nmr.mgh.harvard.edu/fswiki/CorticalParcellation) and subcortical segmentation ‘FreeSurfer_DKT_subcortical’ (surfer.nmr.mgh.harvard.edu/fswiki/FreeSurferVersion3) encompassed coordinates for all the listed regions.

#### Structural alignment to function

The white matter structure facilitating neural regions implicated in cognitive flexibility were mapped using the Human Connectome Project tractography atlas of the young adult white matter 2 mm isotropic template (HCP1065_tractography) (brain.labsolver.org/hcp_template.html). In this work, we refer to this collection of tracts as the structure of cognitive flexibility. The HCP1065 tractography data is a group average constructed from a total of 1,065 subjects, which was acquired using a multishell diffusion scheme with b-values equal to 990, 1,985, and 2,980 s/mm^2^ with 90 diffusion sampling per b-shell. The in-plane resolution and slice thickness were both equal to 1.25 mm. The diffusion-weighted images were resampled at 2.0 mm isotropic during q-space diffeomorphic reconstruction ^26^ to obtain a map of the spin distribution function.^27^ The final white matter template consisted of a 2 mm isotropic map of quantitative anisotropy (QA) values. This template was used to trace the wiring subserving cognitive flexibility function because of its structural coherence with the Neurosynth functional meta-analysis data. Nonetheless, the effects of age/age-group were not evaluated in this diffusion template.

Fibers were reconstructed from the QA template in DSI-Studio (dsi-studio.labsolver.org),^28,29^ using a mean diffusion distance of 1.25 mm, three orientations per fiber, an angular cut-off of 45°, a step size of 1.0 mm, a minimum length of 5 mm, spin density function smoothing of 2 mm, a maximum length of 400 mm, and a QA threshold set to 0.1 (i.e., selection based on the diffusion signal density in the colony-stimulating factor). Tracts with lengths shorter than 5 mm or longer than 400 mm were discarded, and 25 topology pruning iterations were used to trace 10,000,000 streamlines from the template.

We then pursued a parallel approach for the alignment of functional and structural data, as previously reported.^30,31^ Anatomical regions identified in the meta-analysis uniformity map were separated into two files containing cortical or subcortical regions of interest. Each was loaded into DSI-Studio atop the QA template of average adult white matter and dilated to expand the region labels across the grey-white matter boundary. Neural regions in the cortical ribbon were dilated 8 mm, and voxels located in subcortical structures were dilated 4 mm because the gray-white matter interface in subcortical structures is demonstrably thinner. Dilation-filled voxels contained only voxels outside regions of interest. Tracts obtained from the whole brain tractography were converted from endpoints to regions of interest. Streamlines not terminated in the meta-analysis– derived regions of interest were deleted. Finally, the HCP1065 atlas of white matter parcellations was used for recognition and clustering of the remaining streamlines into anatomical brain tracts. The identified brain tracts compose the average structure of white matter implicated in cognitive flexibility function among adult humans.

#### Comparative cohort analyses

To investigate the effects of age on the white matter structure implicated in cognitive flexibility and its relationship to brain function, we computed myelin-related MRI features of white matter density and homogeneity for a subset of the UKBiobank MRI data (n=301), which has a companion functional and behavioral dataset^24^ and a collection of related multi-component functional connectivity attributes previously mapped.^14^ Limited functional, behavioral, and MRI data (T1w, T2-FLAIR, functional MRI) are available for this dataset in OpenNeuro (openneuro.org/datasets/ds003592/versions/1.0.13).

#### Dataset

Our dataset was comprised of discrete samples from 301 healthy younger and older adults who had been characterized by behavioral, self-report, and functional and structural brain MRI assessments. The distribution of age within each age group was tested for normality by computing the kurtosis and skewness over the age variables. The group of younger adults had a mean age equal to 22 ± 3 years (43% male; 57% female), and older adults had a mean age equal to 68 ± 6 years (45% male; 54.2% female; 0.8% non-reported biological sex). Demographic characteristics are described per age group in **Table 1**, and additional descriptions of the original MRI data have been reported previously.^24^

**Table 1.** Sample Characteristics.

| Age group | Age (years) | Biological Sex | Race | Ethnicity |
| --- | --- | --- | --- | --- |
| Younger adults<br>N = 181 | Range: 18-34<br>Mean±SD: 22.6 ± 3.3 | Female: 0.57% | 19.5% Asian<br>8.18% Black or African American<br>4.41% Mixed<br>60.38% White<br>5.03% Other<br>2.5% Not provided | 81.76% Non-Hispanic or Latino<br>10.69% Hispanic or Latino<br>7.55% not provided |
| Older adults<br>N = 120 | Range: 60-89<br>Mean±SD: 68.6 ± 6.4 | Female: 0.55% | 2.54% Asian<br>2.54% Black or African American<br>4.41% Mixed<br>92.38% White<br>2.54% Other | 89.83% Non-Hispanic or Latino<br>1.69% Hispanic or Latino<br>8.48% not provided |

#### Task-based measures related to cognitive flexibility

Set-shifting, inhibitory control, and information updating, which are evaluated through a working memory task, are abilities within the broader cognitive category of executive functioning. In this work, we focused on assessing cognitive flexibility through the primary component of dimensional switching, which is closely related to set-shifting and relies on inhibition and updating. To investigate the associations between white matter features and brain function using behavioral scores, we selected three tasks from the NIH Toolbox (www.nihtoolbox.org) cognitive battery available in the dataset. The dimensional change card sorting, flanker inhibitory control, and list sorting working memory tasks were used. Normalized t-scores computed from task performance are available for the dataset and were used as individual measurements of function.

#### Age effects on functional performance

The distribution of the task score data across the entire age range was evaluated qualitatively (without model), and the outcome plots express distributions within a 95% confidence interval (CI). The distribution of performance scores in each task per age group was tested for inequality using an independent sample *t*-test. Within each age group, the relationship between age and performance in set-shifting, inhibitory control, and working memory tasks was assessed in JASP by linear regression using the forward model y = β_0_ + x.β_1_ + ℇ, where β_0_ is the intercept, β_1_ is the model coefficient, and ℇ is the standard error. The significance level was set at *p* ≤ 0.05, and the influence of sex as a factor was tested. Missing cases were handled listwise.

#### MRI features

MRI features of macromolecular properties of the white matter were computed for all subjects included in the UK Biobank MRI dataset. Anatomical T1w and T2-FLAIR data underwent orientation alignment using the afni (afni.nimh.nih.gov) ‘3dresample – prefix’ function.^32^ T2-FLAIR images were resampled to match spatial resolution with T1w data using the Matlab (www.mathworks.com) ‘imresize3’ function. Volumes were then co-registered using FLIRT, a linear triangulation method in FSL (fsl.fmrib.ox.ac.uk/fsl), skull stripped using the FSL ‘bet2’ function, and masks of brain tissue and non-brain tissue were saved.^33^ Image intensities were standardized using a non-brain-tissue-based equalization method,^34^ which computes intensity corrections using the intensity distribution of stripped tissue (e.g., non-brain tissue). The standardization was then applied to brain tissue voxel-wise using z-score criteria, which are computed with respect to the standard intensity distribution of the ICBM152 anatomical brain atlas, including the cerebellum. Intensity in T2-FLAIR was checked for field bias using standard ghosting criteria.^35^ A point-by-point map of the T1/T2-FLAIR ratio was then computed for each subject.^21^ This ratio was chosen because of its relatively high contrast for segmentation of brain structures in anatomical images.^21^ The HCP1065 anatomical template of the brain was warped to each individual subject’s T1w space using the Matlab ‘imregister’ function, which employs an ‘affine3’ method for volumetric overlay and allows the subject’s anatomy to be kept intact. The transformation matrix was then applied to the atlas’ parcellation mask. An atlas-based parcellation mask of the white matter was then applied to extract properties of 68 anatomical parcels from each subject’s brain (cranial nerves were not considered). In the case of a parcel overlay post affine transformation, voxel membership was selected randomly.

The parcel-specific mean intensity in T1w (*m*) was computed as a measure of comparable macromolecular density, with greater *m* signifying greater macromolecular density along each tract and the counterpart of lower *m* signifying reduced tissue density.^36^ We calculated *m* as the mean value in a distribution of standardized T1w values obtained from voxels within each white matter parcel. Parcel-specific kurtosis (*k*) of the T1/T2-FLAIR ratio was computed as a myelin-related measure of tissue homogeneity throughout each white matter structure. T1/T2-FLAIR ratio has been applied as myelin-related quantity to differentiate myelination levels across different brain structures.^21^ Here, we compute the kurtosis *k* to assess the homogeneity or dispersion of myelin-related signals within each anatomical parcel. The related biological interpretation of *k* is that greater apparent homogeneity is associated with greater kurtosis (e.g., leptokurtic trends) or having most of the data centered around a common value, whereas lower kurtosis (i.e., platykurtic trends) or carrying more data in the tails of the distributions, are interpreted as nonhomogeneous appearance along each brain pathway.^37^ We did not compute *k* for subjects with no T2-FLAIR data. Measures representative of global features of the white matter implicated in cognitive flexibility, *m* and *k*, were also computed for the conglomerate of white matter tracts. We calculated *m* and *k* as the mean and kurtosis of a larger distribution (not tract-specific) containing data from voxels of all 32 implicated bundles. Whole-aggregate *m* and *k* represent the mean T1w (post standardization) and the kurtosis of T1w/T2-FLAIR across the entire white matter structure subserving cognitive flexibility function.

#### Age effects on white matter

The effects of age were evaluated for each white matter parcel (i.e., brain pathway) at an age-group level using independent samples Student’s *t*-test in JASP (jasp-stats.org). Missing values were handled by excluding cases per dependent variable. Levene’s test was performed to confirm the homogeneity of variances, and interpretation was given to probability values *p* > 0.05. Furthermore, the effects of age were also evaluated for each parcel among subjects within each age group using the Bayesian-Pearson correlation method in JASP. ^38^ Pearson’s rho was used to estimate the population correlation coefficient, no valence was assumed, and a Bayes factor of 10 (BF_10_) was chosen for the likelihood boundary.

#### Brain structure and function relationships

Associations between behavioral task performance and MRI features of each brain parcel were examined using a Bayesian-Pearson correlation method in JASP. ^38^ Pearson’s rho was used to estimate the population correlation coefficient, no valence was assumed, and a BF_10_ was chosen for the likelihood boundary.

## Results

### White Matter Structure Implicated in Cognitive Flexibility

The Neurosynth meta-analysis,^19^ which included data from 120 neuroimaging studies and used search terms associated with shifting, identified 5,124 regional neuro-activations that were significantly associated with set-shifting functions within a 95% CI. Synthesized studies included loadings between 0.647 and 0.038, thus representing an inclusionary range of neural variance explained by the included studies. The specificity of the term search was estimated to be 0.885 for the term search when associations with abilities other than shifting were considered with effect sizes >2. These activations largely overlap with regions previously reported as implicated in cognitive flexibility among humans,^2^ including the inferior frontal junction, anterior cingulate cortex, angular gyrus, anterior insula, dorsolateral prefrontal cortex, and inferior parietal lobule.

An atlas-based parcellation analysis revealed that these Neurosynth-derived activations implicated with cognitive flexibility overlapped with 36 neural regions described in four distinct atlases of the human brain, including the posterior and caudal anterior cingulate cortices; superior and middle frontal gyri; and the caudal, superior, middle frontal, and orbitofrontal cortices, in addition to bilateral striatum, putamen, and caudate. The conglomerate of tracts that fed into these neural regions defines the white matter structures subserving cognitive flexibility functions in the human brain. This interconnected wiring is composed of 32 white matter pathways described in the HCP1065 tractography atlas,^20^ which encompasses 82 possible segmentations (68 for bundles and 14 for cranial nerves). The white matter structure of cognitive flexibility includes a few of the longest white matter pathways of the human brain (e.g., cingulum bundle, frontal and parietal aslant tracts, the uncinate and the left arcuate fasciculus, and the inferior, middle, and superior portions of the longitudinal fasciculus). Fig. 2 shows the neural regions implicated in cognitive flexibility and their white matter structures. Additional details are described in Table 2 along with the relevant anatomical atlases.

**Figure 2.**
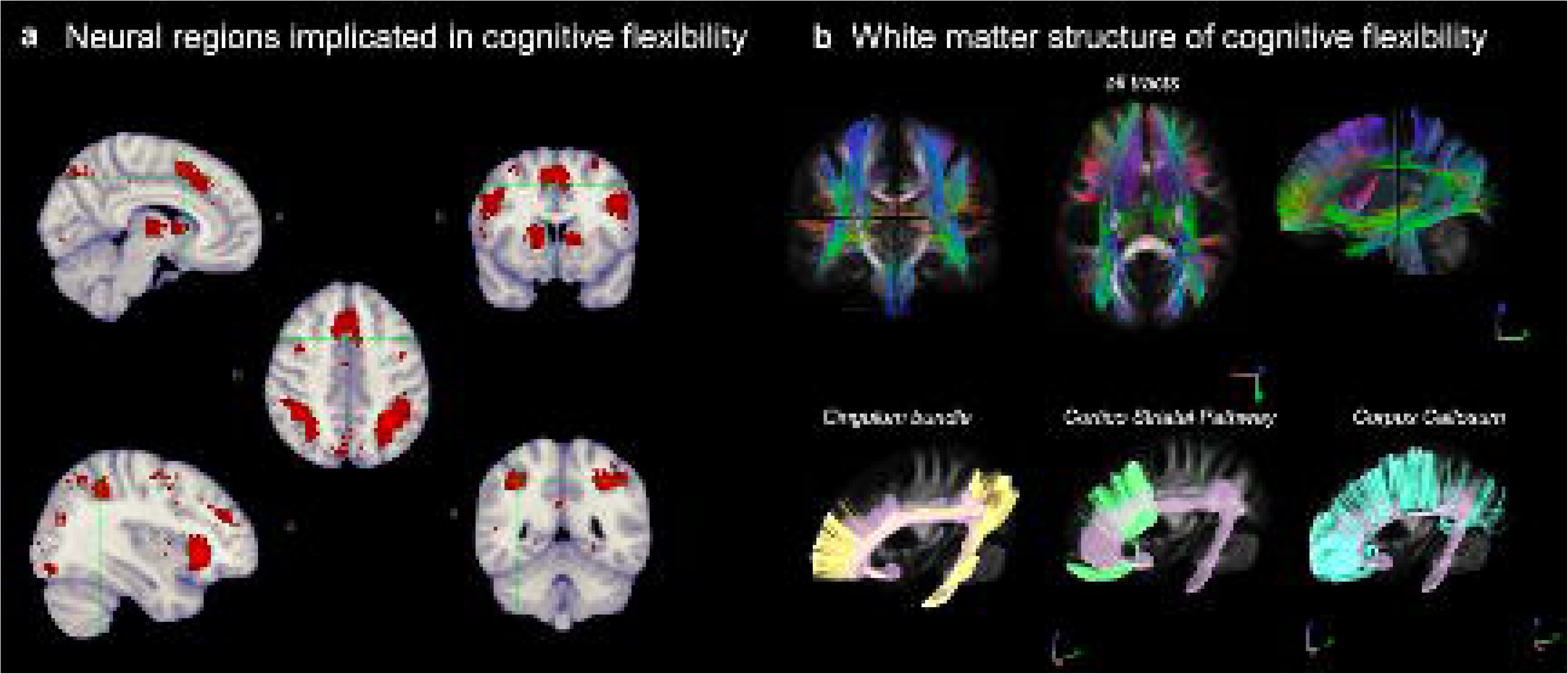
A, Neural regions active during cognitive flexibility tasks (e.g., set-shifting; 95% CI) include the cingulate cortex, several areas of the orbitofrontal and frontal cortex, in addition to the putamen, caudate, and striatum. The complete list of implicated regions derived from the term meta-analysis can be found in ***B***. The underlying white matter structure of the cognitive flexibility function (top row) was found to be composed of 38 independent tracts, which are also listed in **Table 2**. The intricacy of this wiring is illustrated by the common convergency of the left cingulum bundle, the cortico-striatal pathway, and the corpus callosum into distinct regions of the cingulate, prefrontal, and frontal cortex and across the various neural areas of cognitive flexibility. Twenty million tracts were traced by q-space diffeomorphic reconstruction (DSI-studio.labsolver.org)^10^ using the quantitative anisotropy tractography data from the Human Connectome Project atlas of the adult white matter.

**Table 2.**
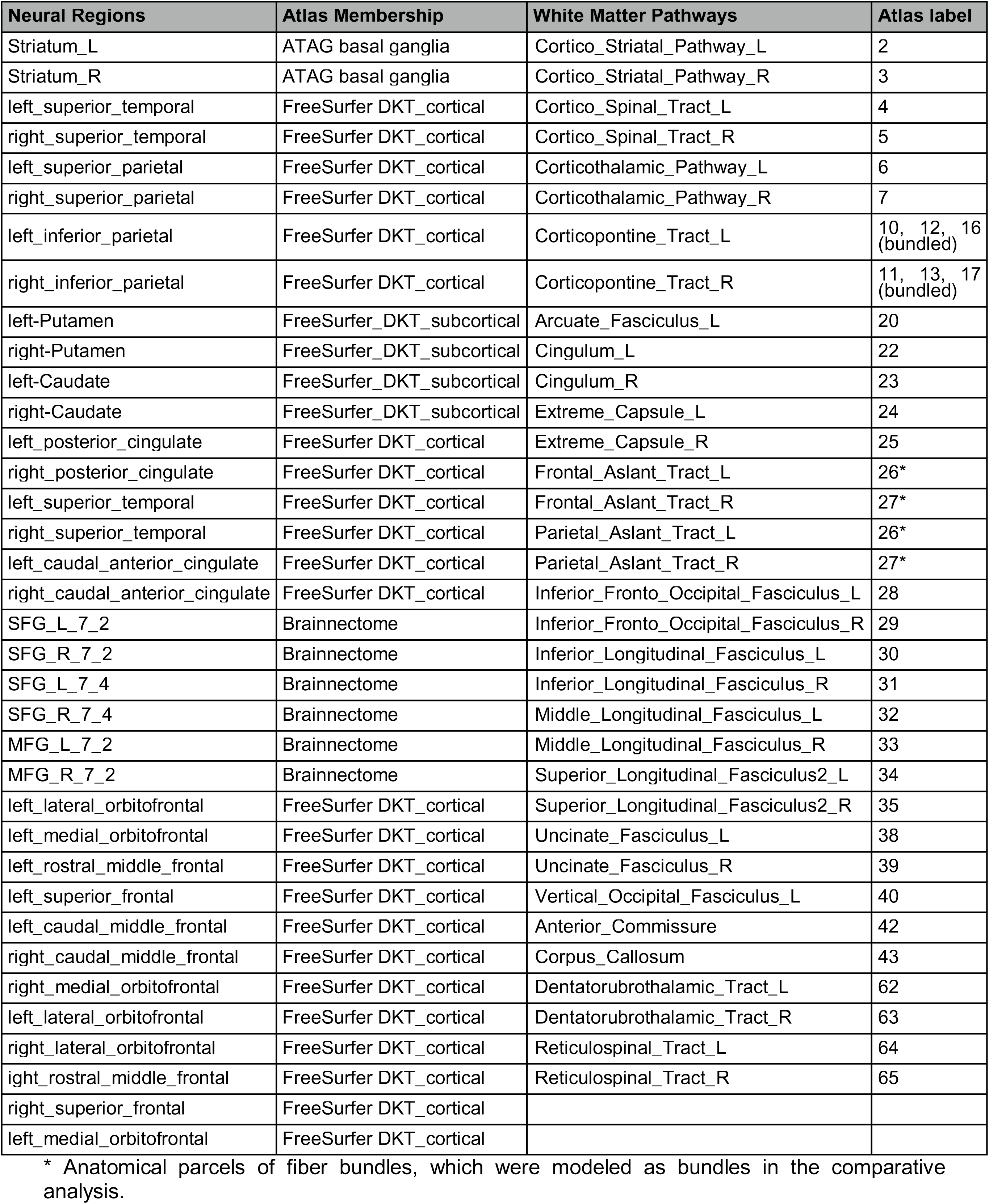
Neural regions of cognitive flexibility and its integrating white matter structure.

| Neural Regions | Atlas Membership | White Matter Pathways | Atlas label |
| --- | --- | --- | --- |
| Striatum_L | ATAG basal ganglia | Cortico_Striatum_Pathway_L | 2 |
| Striatum_R | ATAG basal ganglia | Cortico_Striatum_Pathway_R | 3 |
| left_superior_temporal | FreeSurfer DKT_cortical | Cortico_Spinal_Tract_L | 4 |
| right_superior_temporal | FreeSurfer DKT_cortical | Cortico_Spinal_Tract_R | 5 |
| left_superior_parietal | FreeSurfer DKT_cortical | Corticothalamic_Pathway_L | 6 |
| right_superior_parietal | FreeSurfer DKT_cortical | Corticothalamic_Pathway_R | 7 |
| left_inferior_parietal | FreeSurfer DKT_cortical | Corticopontine_Tract_L | 10, 12, 16<br>(bundled) |
| right_inferior_parietal | FreeSurfer DKT_cortical | Corticopontine_Tract_R | 11, 13, 17<br>(bundled) |
| left-Putamen | FreeSurfer_DKT_subcortical | Arcuate_Fasciculus_L | 20 |
| right-Putamen | FreeSurfer_DKT_subcortical | Cingulum_L | 22 |
| left-Caudate | FreeSurfer_DKT_subcortical | Cingulum_R | 23 |
| right-Caudate | FreeSurfer_DKT_subcortical | Extreme_Capsule_L | 24 |
| left_posterior_cingulate | FreeSurfer DKT_cortical | Extreme_Capsule_R | 25 |
| right_posterior_cingulate | FreeSurfer DKT_cortical | Frontal_Aslant_Tract_L | 26* |
| left_superior_temporal | FreeSurfer DKT_cortical | Frontal_Aslant_Tract_R | 27* |
| right_superior_temporal | FreeSurfer DKT_cortical | Parietal_Aslant_Tract_L | 26* |
| left_caudal_anterior_cingulate | FreeSurfer DKT_cortical | Parietal_Aslant_Tract_R | 27* |
| right_caudal_anterior_cingulate | FreeSurfer DKT_cortical | Inferior_Fronto_Occipital_Fasciculus_L | 28 |
| SFG_L_7_2 | Brainnectome | Inferior_Fronto_Occipital_Fasciculus_R | 29 |
| SFG_R_7_2 | Brainnectome | Inferior_Longitudinal_Fasciculus_L | 30 |
| SFG_L_7_4 | Brainnectome | Inferior_Longitudinal_Fasciculus_R | 31 |
| SFG_R_7_4 | Brainnectome | Middle_Longitudinal_Fasciculus_L | 32 |
| MFG_L_7_2 | Brainnectome | Middle_Longitudinal_Fasciculus_R | 33 |
| MFG_R_7_2 | Brainnectome | Superior_Longitudinal_Fasciculus2_L | 34 |
| left_lateral_orbitofrontal | FreeSurfer DKT_cortical | Superior_Longitudinal_Fasciculus2_R | 35 |
| left_medial_orbitofrontal | FreeSurfer DKT_cortical | Uncinate_Fasciculus_L | 38 |
| left_rostral_middle_frontal | FreeSurfer DKT_cortical | Uncinate_Fasciculus_R | 39 |
| left_superior_frontal | FreeSurfer DKT_cortical | Vertical_Occipital_Fasciculus_L | 40 |
| left_caudal_middle_frontal | FreeSurfer DKT_cortical | Anterior_Commissure | 42 |
| right_caudal_middle_frontal | FreeSurfer DKT_cortical | Corpus_Callosum | 43 |
| right_medial_orbitofrontal | FreeSurfer DKT_cortical | Dentatorubrothalamic_Tract_L | 62 |
| left_lateral_orbitofrontal | FreeSurfer DKT_cortical | Dentatorubrothalamic_Tract_R | 63 |
| right_lateral_orbitofrontal | FreeSurfer DKT_cortical | Reticulospinal_Tract_L | 64 |
| right_rostral_middle_frontal | FreeSurfer DKT_cortical | Reticulospinal_Tract_R | 65 |
| right_superior_frontal | FreeSurfer DKT_cortical |  |  |
| left_medial_orbitofrontal | FreeSurfer DKT_cortical |  |  |

\* Anatomical parcels of fiber bundles, which were modeled as bundles in the comparative analysis.

### Age Effects on Cognitive Flexibility Function

Functional performance in set-shifting, inhibitory control, and working memory tasks significantly differed between age groups (*p* < 0.001) (Fig. 3). Set-shifting abilities appeared to hold an inverse relationship with age, with a logarithmic decay appearance within the 95% CI (Fig. 3*A* left). When evaluated within each age group (younger or older) by linear regression, a significant association was found between set-shifting abilities and age among older adults (β_1_= -0.462, ℇ = 0.118, t = -3.906, *p* < 0.001) but not among younger adults (*p* = 0.859) (Fig. 3A right panels). This result indicates that the decline in the set-shifting function may be particularly relevant in late life.

**Figure 3.**
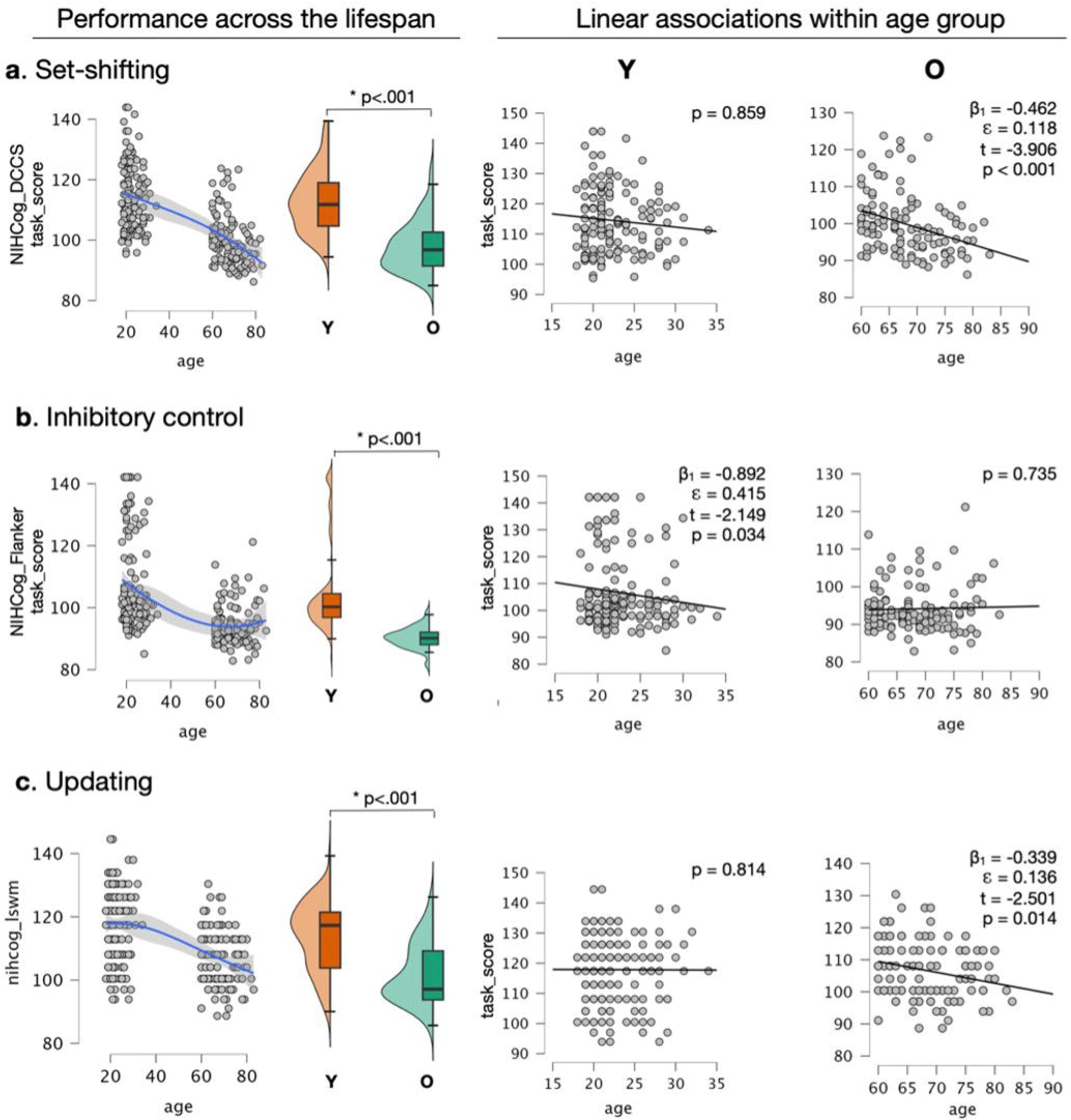
The consonance between brain processes related to cognitive flexibility function and age. ***A*,** Set-shifting abilities exhibited a logarithmic relationship (left panel; 95% CI) across age ranges of earlier and later adulthood, significantly differing between groups (center panel box plots; *p* < 0.001). Among older adults, set-shifting exhibited a significant linear relationship with age (β_1_ = -0.462, ℇ = 0.118, t = -3.906, *p* < 0.001) but not among younger adults (*p* = 0.859). ***B*,** Inhibitory control abilities exhibited a hammock-shaped relationship (left panel, 95% CI) across the same age ranges, similarly differing between the older and younger age groups (center panel box plots; p<0.001). A linear association between age and inhibitory control function was found only among younger adults (β_1_ = -0.892, ℇ = 0.415, t = -2.149, *p* = 0.034) but not among older adults (*p* = 0.735). ***C*,** Working memory abilities exhibited overall behavior similar to set-shifting, varying significantly between age groups (left panel; p<0.001) and presenting an inverse linear association with age among older adults (β_1_ = -0. 339, ℇ = 0.136, t = -2.501, *p* = 0.014), but not among younger adults (*p* = 0.814). These results demonstrate the nature of function-specific associations between biological age and human abilities, further emphasizing intrinsic relationships that exist across the age strata.

On the other hand, inhibitory control exhibited a non-linear relationship with age within the 95% CI (Fig. 3*B* left), with a significant linear association with age among younger adults (β_1_= -0.892, ℇ = 0.415, t = -2.149, *p* = 0.034), but non-significant curvilinear association among older adults (p = 0.735) (Fig. 3*B* right panels). This result indicates that inhibitory control abilities are likely invariant in later life.

Similar to set-shifting, working memory abilities (information updating) exhibited a logarithmic decay with age within the 95% CI (Fig. 3*C* left), with a significant linear decay among older adults (β_1_= -0. 339, ℇ = 0.136, t = -2.501, *p* = 0.014), but not among younger adults (*p* = 0.814) (Fig. 3*C* right panels). This result indicates that information-updating abilities, such as working memory, decline as a function of age in later life.

### Age Effects on the White Matter Structure of Cognitive Flexibility

The macrostructural characteristics of each relevant tract were analyzed across younger and older adult groups, focusing on two main parameters: (1) increased mean in standardized T1w (*m*), and (2) tail trends in distributions of myelin-related appearance, as indicated by the kurtosis of T1/T2-FLAIR ratios (*k*) (see Fig. 4*A-C*).^21^ Elevated *m* values suggest higher macromolecular density within the tract, while lower *m* values indicate reduced tissue density. The interpretation of *k* revolves around the apparent homogeneity within the tract: higher kurtosis (leptokurtic trends) suggests greater uniformity with most data clustered around a central value, whereas lower kurtosis (platykurtic trends) implies a non-uniform appearance with data distributed towards the tails.

**Figure 4.**
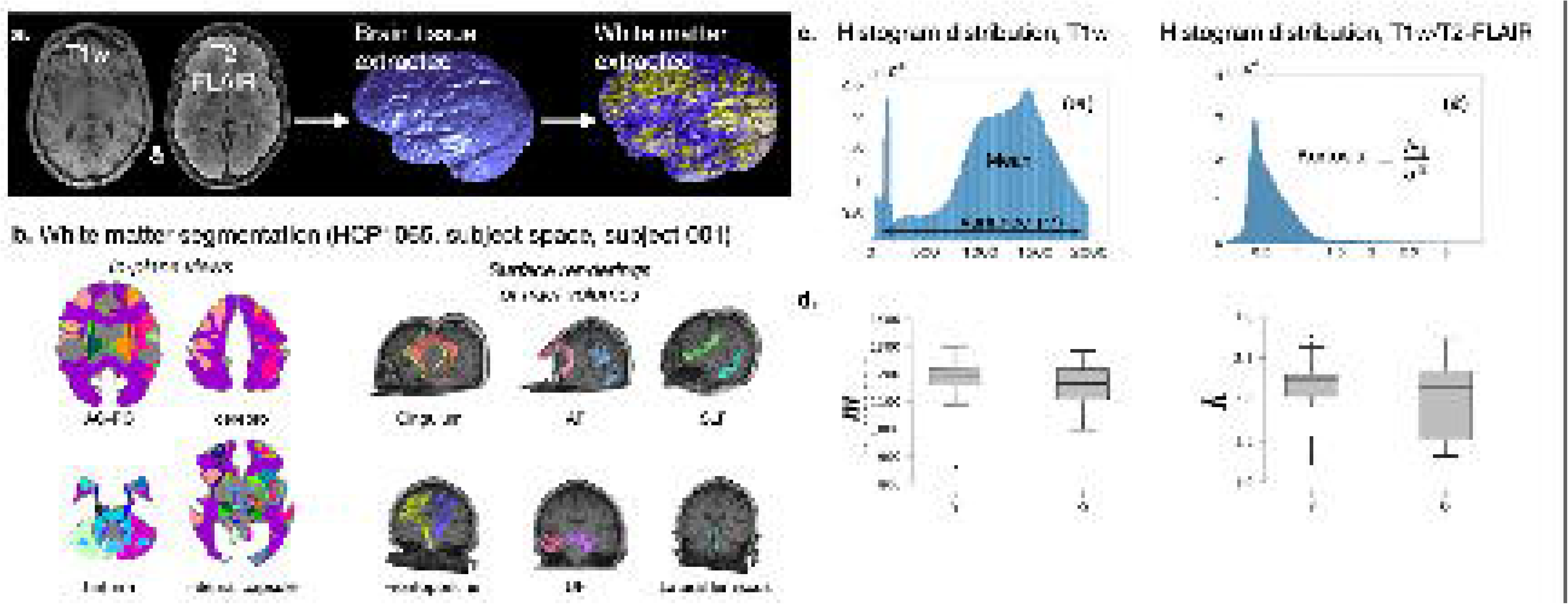
A, Extracted brain white matter showing the representativeness of the encased volume. **B,** Region segmentation of white matter in the subject’s native space showing the HCP1065 atlas-based parcels in consecutive axial planes (top) and their surface rendering (bottom). **C,** Distributions of image-derived parameters of macrostructural integrity for a representative case. *m*: region-specific T1w mean; *k* region-specific kurtosis of the T1w/T2-FLAIR ratio. **D,** As a whole, the homogeneity of the white matter structure of cognitive flexibility significantly differs among younger and older adults—*m*: t(30)=9.26, *p* < 0.001*; k*: t(27)=3.24, *p* = 0.003.

As a whole aggregate, the white matter structure implicated in cognitive flexibility exhibited significant age-group differences in both *m* and *k* (*m*: t[30] = 9.26, *p* < 0.001; *k*: t[27] = 3.24, *p* = 0.003). These results indicate that, on average, the white matter structure of cognitive flexibility is significantly more dense and more homogenous among younger adults (higher *m*, leptokurtic trend in *k*) than among older adults (lower *m*, platykurtic trend in *k*). The distribution of both variables, the structure’s phenotypical MRI appearance (i.e., *m*), and the trends of myelin-related homogeneity (i.e., *k*) are shown in Fig. 4*D*. The statistical dispersion of these variables qualitatively depicts within-group age effects, and indicates that, on average, the density and homogeneity of the entire structure of cognitive flexibility declines in the later decades of life.

Table 3 shows quantification of the age-group effects for each white matter tract in the structure of cognitive flexibility. Although the entirety of the structure of cognitive flexibility carries significant differences with age group membership among adults, once parceled out, not all segmented pathways carried significant differences in density or homogeneity between age groups. In particular, age-group-related variations in tract homogeneity were not detected in the left frontal and parietal aslant tracts, right inferior fronto-occipital fasciculus, left inferior longitudinal fasciculus, left dentatorubrothalamic, and right reticulospinal tracts. These findings indicate that, on average, the overall tissue homogeneity across the structure subserving cognitive flexibility is significantly different within those two periods of human life, but age-group-specific variations are selective to specific brain pathways (stars in Table 3; 95% CI), with several white matter bundles which are also implicated in sensory, motor, learning, and sensory processing not displaying detectable differences across age groups.

**Table 3.**
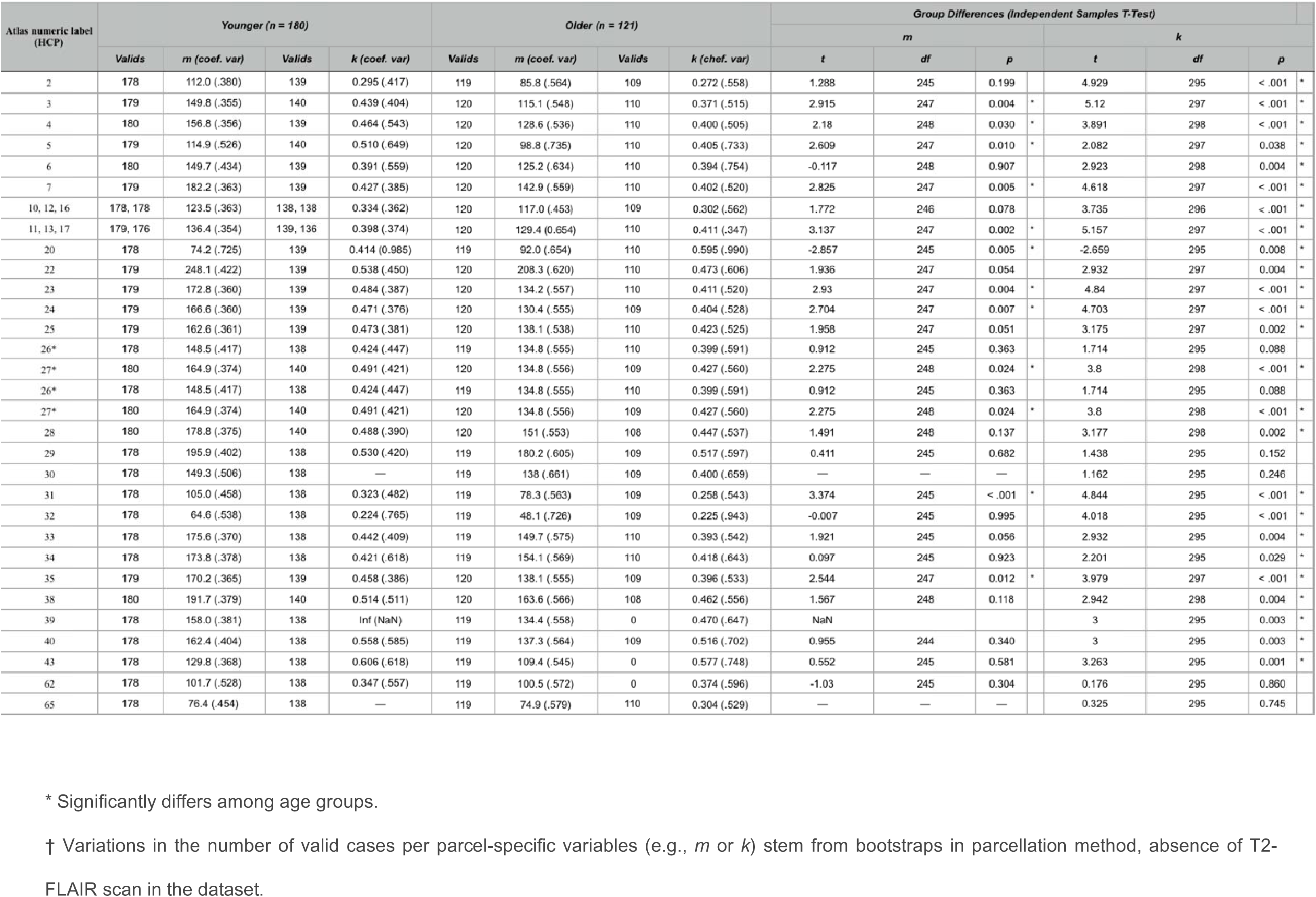
Age-group effects on myelin-related homogeneity per white matter tract in the structure of cognitive

Additionally, a significant age effect on the entirety of the white matter structure was observed among older but not among younger adults. The myelin-related homogeneity *k* was found to be inversely correlated with chronological age among older adults (*r* = -0.643, *p* < 0.001, 95% CI). This indicates that the white matter integrity of these aggregated brain structures decreases with increasing age later in life. This relationship remained nearly unchanged when biological sex was considered (*r* = -0.596 [males] and r = -0.604 [females], *p* < 0.001 [both]).

### Relationship between brain function and the structure of cognitive flexibility

Three distinct functions related to cognitive flexibility were considered: dimensional set-shifting, inhibitory control, and working memory.^1^ The association between brain function and the white matter characteristics of the structure implicated in cognitive flexibility was evaluated for each brain pathway using Bayesian-Pearson correlations reported at BF_10_>100 and 95% CI. These relationships were evaluated within age groups. Fig. 5 shows the white matter pathways that were found to present function-correlated appearance within each life period colored by functional construct (singular constructs or shared among multiple constructs). Tract-selective associations are detailed below for each age group.

**Figure 5.**
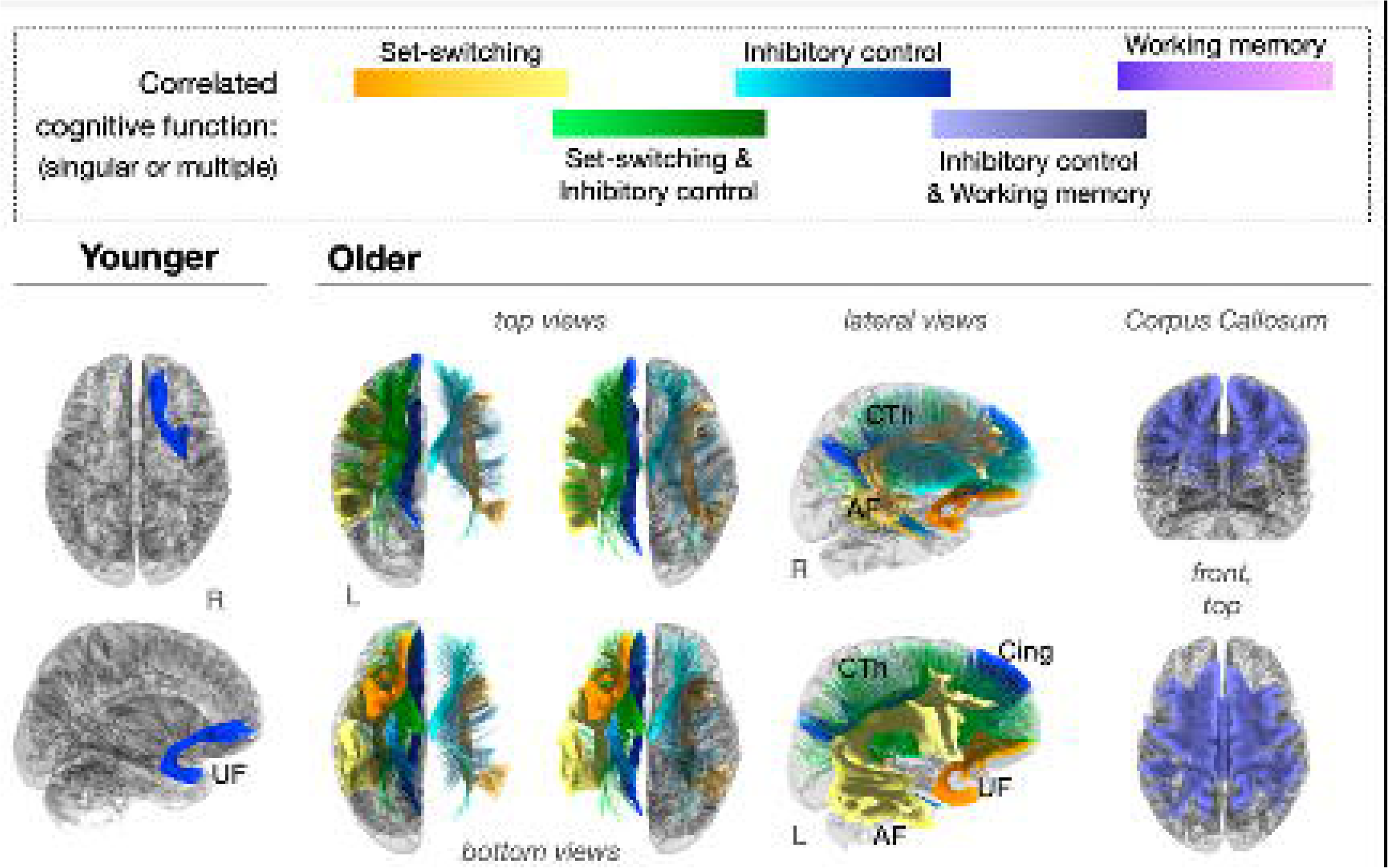
Age effects on brain tracts that present function-correlated appearance in MRI. A subcollection of brain tracts among those with membership in the circuit of cognitive flexibility show significant correlations between white matter homogeneity and functional performance in older adults but not in younger adults. This finding indicates that age-related changes in cognitive flexibility function are reflected in the integrity of specific white matter compartments. Tract colors indicate the correlated cognitive function. AF, arcuate fasciculus; UF, uncinate fasciculus; CTh, corticothalamic projections; Cing, cingulum bundle.

Among younger adults, inhibitory control (e.g., t-score of performance in the flanker inhibitory control task) was the only domain that exhibited a significant direct association with white matter integrity. Specifically, *m* computed for the right uncinate fasciculus (UF) inversely correlated with inhibition ability (*m_UF_*right: r = -0.202, *p* = 0.043). Tract homogeneity (i.e., *k* values) was not significantly correlated with task performance among younger adults. These results indicate that variability in white matter structure that exists in earlier adulthood was not a significant explanatory variable of functional performance.

Among older adults, several distinct tracts carried significant correlations with brain functions. Performance in a set-shifting task was observed to negatively correlate with the appearance of the arcuate fasciculus (AF) bilaterally (*m_AF_* right: r = -0.487, *p* < 0.001; *m_AF_*left: r = -0.488, *p* < 0.001) and a unilateral, negative correlation was found between dimensional set-switching function and the left UF (*k_UF_* left: r = -0.221, *p* = 0.003). Additionally, the homogeneity of the corticothalamic projection fibers (Cth) was found to carry a significant left-lateralized correlation with the dimensional set-switching function (*m_Cth_* left: r = -0.119, *p* = 0.048). Given the large degree of connectivity of the cingulum bundle, a white matter pathway that extends throughout all cerebral lobes, its white matter features were expected to correlate with flexibility function. However, curiously, performance in the set-switching task did not correlate significantly with the homogeneity of the cingulum bundle (Cing) in isolation (*k*_Cing_ left: BF_10_ = 0.776; *k*_Cing_ right: BF_10_ = 0.120). Furthermore, significant correlations with functional scores were not observed for the right UF (*m*: BF_10_ = 1.159, *k*: BF_10_ = 0.191), nor for the CC (*m*: BF_10_ = 1.269, *k*: BF_10_ = 0.199) among older adults. These findings indicate that age effects that occur later in life are selective to specific tracts and affect one’s dimensional set-switching abilities.

Performance in an inhibitory control task was found to be significantly correlated with the homogeneity of the association fibers of the corpus callosum (CC) in older adults (*m_CC_*: r = -0.465, *p* < 0.001) and bilaterally with the ChT projections (*m_Cth_* right: r = -0.477, p < 0.001; left: r = -0.479, *p* < 0.001). Additionally, a significant left-lateralized association was observed between inhibitory control and an increase (positive correlation) in homogeneity across the left cingulum bundle among older adults (*k_Cing_* left: r = 0.157, *p* = 0.011). These findings indicate that, in later life, inhibitory control function finds broader, decentralized correlates with brain structure, including both hemispheres amply.

A single significant correlation with white matter integrity was observed for working memory abilities among older adults. Performance in a working memory task was positively correlated with the homogeneity of CC fibers among older adults (*k_CC_*: r = 0.172, *p* = 0.007). This result indicates an unspecific, but rather global, link exists between information updating abilities, such as working memory, and the brain’s white matter in later life.

## Discussion

Our findings support the life-period-related differentiation of white matter tracts implicated in cognitive flexibility as a natural substrate of brain function. We found that (1) the wiring of cognitive flexibility is defined by a collection of specialized tracts, which present undifferentiated characteristics early in adulthood and significantly differentiated features in later life; (2) these macromolecular, myelin-related white matter properties are correlated with individual subprocesses of cognition which are intimately related to the latent construct of cognitive flexibility function (i.e., dimensional switch, inhibition, updating)^1^; and (3) in the later decades of life, the myelin-related homogeneity of specific white matter tracts implicated in cognitive flexibility declines as a function of chronological age.

In this study, we used a novel data-driven approach to segment the average white matter structure subserving brain regions of cognitive flexibility using a white matter template of the human brain (i.e., the HCP1065 tractography-based atlas). Figure 1 presents a diagram of the method, and the primary code used in this analysis is available as an open resource (github.com). Through a meta-analysis in Neurosynth (neurosynth.org), we defined the neural regions reportedly implicated in cognitive flexibility tasks and then traced the structure wiring these brain regions using the HCP1065 template of quantitative anisotropy in white matter. This approach allowed us to analyze brain regions and tracts of specific interest to cognitive flexibility function while evaluating age effects and associations with functional domains. Furthermore, this method allowed us to evaluate age effects across the entirety of the delineated structure and in a tract-selective manner. Importantly, we present data that emphasizes the value of evaluating white matter structures in the context of specific brain functions.

White matter features were investigated in terms of MRI-derived quantities to assess macromolecular tissue properties (T1w, T2-FLAIR). Such approaches center the work on the consideration of age- and function-related variations in homogeneity within white matter bundles rather than structural alignment or coherence, which can be assessed by diffusion-based measures (e.g., fraction of anisotropy, radial diffusivity). Commonly, radiologic evaluations of these MRI modalities point to hypo- or hyperintensities as degenerative features, and the appearance of white matter hyperintensities in T2-FLAIR is observed in pathology and brain injury.^22^ Several studies focused on the coherence of white matter axons, which is a parameter measured by diffusion MRI, document correlations between white matter thinning (reduced tissue coherence, reduced fraction of anisotropy [FA]) and neuropathology or decline in functional scores. Specifically in the context of cognitive flexibility, in vivo diffusion studies have documented that changes in parameters of diffusivity within the white matter, such as myelin-related radial diffusivity and fraction of anisotropy, are correlated with advanced age in healthy aging adults.^23^ These changes in global and tract-specific incoherence of white matter have been cited as the origin of concomitant functional decline.^13^ Although intriguingly parallel to the knowledge of (micro)structural features of age-related pathology that appear in the cerebral tissue of some older adults, these findings are restricted by the limitations of diffusion imaging measures. It remains unclear whether diffusion MRI can distinguish between features of natural white matter differentiation and pathology. The data presented in this work considers associations between macromolecular tissue features, cognitive flexibility function, and age.

The MRI dataset used in this work was acquired in adults with no documented pathology or cognitive deficits, and the variability among the sample is attributed to natural age.^24^ We evaluated the homogeneity of the white matter bundles in terms of the asymmetry of the T1w/T2-FLAIR ratio distribution, therefore encompassing tissue irregularities that may exist due to (natural age-related) hypo- or hyperintensities along the bundles. The volume of white matter hyperintensity was not considered explicitly, and the consequent limitations are discussed. In this work, hyperintensities were considered implicitly through the measure of kurtosis of the T1w/T2-FLAIR distribution, *k*.

The process of natural aging is defined in the context of neurosciences as the time-related deterioration of structural and functional cellular processes, which results in the maturation of brain tissues.^25^ Peters and Folger have documented the remarkable resemblance between age- and pathology-related features of neuronal bodies and myelin stacks, emphasizing the relevance of function-centered investigations in aging studies. Age-related changes in neural structure and function overlap with features of late-life pathophysiology and may manifest at different degrees across the lifespan in the presence of other risk factors. This poses an added challenge to studies on the effects of age on brain structure and function. In this work, we propose that by deriving white matter structures related to cognitive flexibility from meta-analysis-derived functional patterns, we reveal white matter structures representative of population-level structure-function associations (see Fig. 1, left panel). We further applied these structures to an independent dataset (the UK Biobank subset) that includes MRI and functional/behavioral data from adults with no impairment or pathology (see Fig. 1, right panel). Lastly, we derived associations and the effects of age. Although this methodological approach has limited specificity to pathological processes, it reveals age-related macromolecular features of white matter directly associated with cognitive flexibility function and the effects of age.

The homogeneity across the entirety of the resulting white matter structure was found to differ between early (e.g., <45 years) and later adulthood (e.g., >65 years). However, not all tracts within the structure showed significant age-group differences (95% CI), as shown in Table 2. These findings were computed after testing for equality of variances per tract (i.e., Lavene’s *p* > 0.05) and highlight that the white matter structure of cognitive flexibility differs in different periods of adult life, with older adults carrying a significantly less dense and homogeneous type across select tracts and appearing more left-lateralized (see Fig. 5).

The correlation between white matter homogeneity and function also varies across different periods of life. We found that among younger adults, the density and homogeneity of specialized tracts are not associated with the network-level functions of dimensional switching or with updating, but an inverse association exists between the appearance of the right uncinate fasciculus and inhibitory control. On the other hand, among older adults, a left-lateralized pattern involving a few projection, association, and commissural fibers was found to correlate with the specific functions. These findings agree with the elevated functional network manifold eccentricity (i.e., a graph measure of how diffuse or decentralized networks are) found by Setton et al. 2023 in the same dataset.^14^ Using multi-echo resting state BOLD series associated with this dataset,^1^ the study found that canonical functional networks that are related to cognitive flexibility (e.g., default-mode, dorsal attention, fronto-parietal) are physiologically more integrated among older adults, with particular components gaining greater functional specialization later in life. Our findings on the age-related specialization of these same subjects’ white matter further corroborate an argument for a fundamental myelin-related mechanism of brain aging that may exist in synchrony with functional aging.

The patterns of white matter aging and their relationship to function have been charted across the lifespan.^2,16^ Grydeland et al. have shown that white matter ages differently in different areas of the brain, presenting either an inverted U-shaped dependence with chronological age or a long-lasting linear and positive relationship with aging (indicating myelination increases in some areas as one ages). Both these association types go hand-in-hand with cognitive performance.^3,5,6^ Here, we show that age effects also exist at the system-structure level. For the structure of cognitive flexibility, it appears that the orchestrated differentiation of white matter tracts in the later periods of life describes structural regionalization and its consequent functional performance.

### Limitations

A relevant limitation of our analysis is that these correlations were observed considering data collected in normative subjects and thus have limited extension to the interpretation of clinical samples with deficits in cognitive flexibility. Moreover, we investigated age effects across age groups and the effects of age within age groups using a cross-sectional sample. Therefore, these findings have limited sensitivity to individual variability in age-related features and their association with brain function. Additional studies are necessary to determine if the slope of decline in myelin-related homogeneity is an effective predictor of functional impairment or risk and to further understand if tract-selective associations can explain functional deficits.

The fraction of radiologically visible white matter hyperintensity that is present within each white matter bundle was not explicitly considered in this work, thus limiting the specificity of our findings. Given that hypo- or hyperintensities modify the skewness of the distribution of the T1w/T2-FLAIR ratio, our findings are linked to white matter normative age-related features that appear at the population-level in asymptomatic adults. Additional studies focused on measures specific to pathology are necessary to investigate the role of macromolecular hyperintensities in age-related decline.

A limited evaluation of cognitions was conducted in this study. Cognitive flexibility, as measured by set-shifting and two related cognitions, inhibition and updating, was considered. Although a significant age-related, tract-specific distinction was found for the link between white matter homogeneity and function among these cognitions, a larger-scale investigation would be necessary to identify if this function-related differentiation of white matter is a generalized phenomenon across other functions.

Additionally, it is possible that the functional patterns of cognitive flexibility among older adults may significantly differ from that of younger adults, for instance, in the context of age-related reorganization. In this work, meta-analysis was employed to derive the population-level functional pattern of interest, but it is possible that an intrinsic age-related difference exists between the regions of interest for cognitive flexibility among older versus younger adults. Additional studies on age-related changes in functional topology, in combination with diffusion MRI, performed at the individual level are necessary to investigate this limitation.

Furthermore, our study was limited to the evaluation of white matter tracts that can be reliably segmented in anatomical images of the brain. For example, the anterior commissure was identified in the tractography as part of the average white matter structure of cognitive flexibility in humans. However, its segmentation was not feasible using anatomical images of the dataset due to poor voxel resolution (i.e., 1-2 mm), and therefore, its role and age effects remain elusive. Here, we used a limited sample of the UK Biobank MRI dataset, granting that replicability of our findings in other samples is necessary. Analysis of additional datasets may enrich the understanding of age effects on macromolecular features of white matter that are related to cognitive flexibility. Additional studies, including delineation of the brain’s fine tractography performed at the individual level, have the potential to substantiate the understanding of the specific roles that white matter tracts play in sustaining and changing cognitive flexibility function.

## Data, Materials, and Software Availability

All code used for analysis is available on GitHub and can be found at this link: https://github.com/TatianaWolfe/NeuroAge_v5. Subjects’ original images and functional and behavioral data are available in OpenNeuro and can be found at this link: 200219.

## Acknowledgments

The authors acknowledge the technical contributions of Natalie Morris and Deborah Hodges. This work made use of the high-performance computing resource in the Brain Imaging Research Center (BIRC) of the Psychiatric Research Institute at the University of Arkansas for Medical Sciences. NIH grant 5P20GM109005-09 RPL (T.W.).

## Author Contributions

TW: conceived and designed the analysis; created tools for analyses; performed the analysis; wrote the paper. AG: performed the analysis; designed the analysis; MC: designed the analysis; wrote the paper. AP: performed the analysis. LI: designed the analysis; wrote the paper. CK: conceived and designed the analysis. TK: performed the analysis; wrote the paper. GAJ: designed the analysis; wrote the paper.

## Classification

Major classification: Biological Sciences. Minor classification: Neuroscience.

